# Stag2 dependent chromatin remodeling enforces the erythroid-specific Gata1 cistrome

**DOI:** 10.1101/2025.09.15.676332

**Authors:** Varun S. Sudunagunta, Edna M. Stewart, Yi Chen, Hongxia Yan, Sebastian Fernando, Viviana Scoca, Jane J. Xu, Narla Mohandas, Aaron D. Viny

## Abstract

The transcription factor GATA1 has pleiotropic hematopoietic functions, particularly in erythroid and megakaryocytic ontogeny. While mechanistic investigations have uncovered many facets of GATA1 biology, how GATA1 co-regulates divergent cell fates remains only partially characterized. We previously described that loss of Stag2, a member of the cohesin complex and a recurrent mutational target in myelodysplastic syndrome and Down Syndrome associated acute megakaryoblastic leukemia, results in altered chromatin accessibility, transcription factor function, and cell differentiation. Hence, we hypothesized that chromatin accessibility facilitates lineage specificity of GATA1, thereby permitting efficient cellular differentiation. To understand the connection between chromatin accessibility and GATA1, we performed comprehensive studies of erythropoiesis in *Stag2^Δ^* mice. Defects in Stag2-deficient hematopoiesis included reduced numbers of erythroid progenitors (EryPs), impaired terminal erythroid maturation, increased number of MkPs, and increased megakaryocytes. RNA- and ATAC-sequencing of EryPs revealed altered patterns of Gata1 target gene expression with altered accessibility in conjunction with loss of expression of erythroid targets and gain of megakaryocyte targets. Gata1 occupancy was lost at erythroid targets, while occupancy increased at megakaryocyte targets. Functionally, we observed that Stag2-deficient EryPs have diminished erythroid output and augmented megakaryocyte output in orthogonal differentiation assays. Human models and primary MDS patients recapitulated the essential phenotypic and molecular features of our in vivo murine MDS model. Collectively, this study advances the understanding of the interplay between TF function and chromatin accessibility. Moreover, these data suggest a novel conceptual paradigm of dyserythropoiesis in MDS.

**Key Points:** xxxxx

## Introduction

Erythropoiesis is a tightly orchestrated process that transforms multipotent hematopoietic stem cells into mature erythrocytes through successive stages of lineage restriction, nuclear condensation, global transcriptional silencing, and enucleation^1,2^. This process is not only highly efficient – producing over 200 billion red blood cells daily under steady state – but also remarkably adaptable under stress, capable of increasing red cell production 20-fold. Central to the regulation of erythropoiesis is a complex network of epigenetic mechanisms at multiple levels that govern chromatin accessibility, transcriptional dynamics, and nuclear architecture^3–6^. These processes are mediated by both intrinsic and extrinsic factors such as cytoskeletal elements^7,8^, histone modifications^9,10^, vesicle trafficking^11^, and cell adhesion molecules^12^. Together, they ensure balanced hematopoietic output and reserve capacity during stress.

- Stag2 facilitates lineage specificity of Gata1 by modulating chromatin accessibility, permitting balanced and efficient erythropoiesis.
- STAG2 loss in human cells and in MDS patient samples conserves key features of dyserythropoiesis and cistrome dysregulation.

While key erythroid transcriptional networks, hemoglobin switching, and immunophenotypic transitions have been well characterized^10^, the mechanisms that confer lineage specificity remain incompletely understood. Transcription factors (TF) such as GATA1 are broadly expressed across hematopoietic lineages, yet direct distinct differentiation programs in different cellular contexts. For example, GATA1 is expressed in a variety of stem and progenitor populations, yet is also critical in lineage-specific erythroid^13,14^, megakaryocytic^15^, eosinophilic^16^, and mast cell^17^ differentiation. This raises fundamental questions underlying how lineage specificity is achieved. Although cofactors like FOG1 contribute to specificity, emerging evidence suggests that TFs likely also rely on epigenetic context – such as chromatin accessibility and remodeling complexes – to enforce lineage-restricted cell-context specific gene expression programs and ultimately mutually exclusive cell fates^18–20^.

We previously demonstrated that loss of the cohesin subunit STAG2, which is mutated in approximately 10% of MDS, disrupts chromatin accessibility, TF activity, and lineage differentiation^21,22^. Notably, while Stag2 deficiency in lymphopoiesis impairs the opening of lineage-specific chromatin regions, erythropoiesis is uniquely characterized by a failure to close chromatin, suggesting a distinct regulatory mechanism. We hypothesized that defective formation of repressive heterochromatin may impair erythroid differentiation through disruption of lineage-specific transcriptional programs. To validate this hypothesis, we leveraged our transgenic *Stag2^Δ^* mouse model to examine how altered chromatin accessibility influences erythroid TF function and transcriptional output. By profiling chromatin dynamics across erythroid lineage commitment and terminal differentiation, we aimed to define how chromatin remodeling enforces erythroid identity and how its disruption may contribute to disease.

## Methods (also supplementary methods)

### Mice

All animals involved in this study were housed at Columbia University Irving Medical Center (CUIMC) and were utilized for experimental purposes in accordance with approved Institutional Animal Care and Use Committee (IACUC) protocols and animal welfare standards. Mx1-Cre Stag2^fl/fl^ mice were previously described^22^. Recombination of Stag2 was induced by 4 intraperitoneal doses of polyinosinic:polycytidylic acid (pIpC; 20 mg/kg) starting at 6 weeks of age. Terminal endpoints for all mice were no earlier than 4 weeks after the last dose of pIpC. Stag2 excision was verified at the time of terminal analyses as previously described^22^. Mice were assigned to experiments agnostic of sex.

### MDS Patients

Diagnostic bone marrow hematopathology reports from patients with MDS who were treated at CUIMC were analyzed for descriptors of erythrodysplasia. For a subset of patients, digital peripheral blood smear images were analyzed (CellaVision). Patient characteristics are described in supplemental table 1. This study was approved by the Institutional Review Board at CUIMC (AAAU4817, AAAU8073).

### Murine liquid culture

5000 EryPs were sorted into 100 µL of FACS buffer, washed once with PBS, and resuspended in 200 µL of StemSpan^TM^ SFEM II (StemCell Technologies, #09655) supplemented with 1% penicillin/streptomycin, rm-stem cell factor (100 ng/mL; Peprotech), and rh-erythropoietin (4 U/mL; Peprotech)^23,24^. After 48 hours, cells were collected and analyzed by flow cytometry for absolute counts and surface markers .

### Murine colony forming assays

20,000 (erythroid-directing; M3334) or 10,000 (pan-myeloid; M3434) EryPs were seeded in 1 mL MethoCult (StemCell Technologies) supplemented with 1% penicillin/streptomycin and no additional supplemental cytokines in duplicate. Colonies were scored after 48 hours (M3334) via brightfield microscopy (Nikon Eclipse Ts2) and immunofluorescence or on day 8 (M3434) via flow cytometry.

### Human CD34^+^ HSPC liquid culture and shRNA knockdown of STAG2

CD34^+^ cells were isolated from umbilical cord blood, transduced with shRNA, and cultured as previously described^25^. Cells were collected on D5-7, D11, and D15 of culture, stained as previously described, and analyzed by flow cytometry for cell surface markers^26,27^. Additional details are provided in supplemental methods.

### RNA sequencing data analysis

The normalized count matrix was downloaded from GSE138006 and analyzed as per the supplemental methods.

### scATAC sequencing

HSPC (Lineage negative) and Ter119^+^ cells were sorted, sequenced, and analyzed as described in supplemental methods.

### Gata1 CUT&RUN

Cleavage Under Targets and Release Using Nuclease (CUT&RUN) was performed using CUTANA^TM^ ChIC/CUT&RUN Kit and CUT&RUN Library Prep Kit (EpiCypher) as per manufacturer’s instructions. Details and analytic tools used are provided in the supplemental methods.

### Statistical Analysis

All experiments were reported as indicated: *N* or the number of data points indicates the number of independent biological replicates. Statistical significance was evaluated using GraphPad Prism (v.10). Data distribution was assumed to be normal, but was not formally tested. Outliers were not excluded unless indicated.

### Data Availability

Processed sequencing data is made available via NCBI Gene-Expression Omnibus (GEO) at GSE301108. MDS patient derived CD34^+^ bulk RNA-seq was obtained from GSE114922. Reference ChIP-seq experiments were obtained from the ENCODE Project (Gata1 G1ER ENCFF806QBE, Gata1 Megakaryocyte ENCFF366HNL, Tal1 G1ER ENCFF406CDH). Klf1 ChIP-seq from murine fetal liver was obtained from GSE210779^28^ All data generated in this study are available within this manuscript and its supplemental files, or on written request from the corresponding author. Code used for data analysis is available upon written request to the corresponding author.

## Results

### Stag2 coordinates chromatin accessibility during erythroid lineage commitment

To map the dynamic changes in chromatin accessibility throughout early erythroid fate commitment, we performed scATAC sequencing on non-lineage committed hematopoietic stem and progenitor cells (HSPCs) from wild type and Stag2-deficient bone marrow, henceforth referred to as *Stag2^WT^*and *Stag2^Δ^* (**Figure 1A**). Using genes known to be involved in various hematopoietic cell stages, we identified each of the major hematopoietic lineages and mapped each lineage’s differentiation trajectory **(Figure S1A).** Subsequently, we filtered the captured cells for those along the differentiation trajectory for erythropoiesis, retaining cell stages from hematopoietic stem and progenitor cells through the bipotent megakaryocyte/erythroid progenitor (MEP) stage, and into the lineage committed erythroid progenitors (EryP) based on ATAC peak profiling (**Figure 1B; S1B)**. Consistent with prior work, we observed that *Stag2^Δ^* HSPCs have defective erythroid commitment and differentiation (**Figure 1B-C**)^22^. Moreover, our prior transcriptomic analyses identified a delayed erythroid pseudotime, which was evident in progenitor populations by scATAC-seq based on the proportions of inferred cell types (**Figure 1C**).

**Figure 1.**
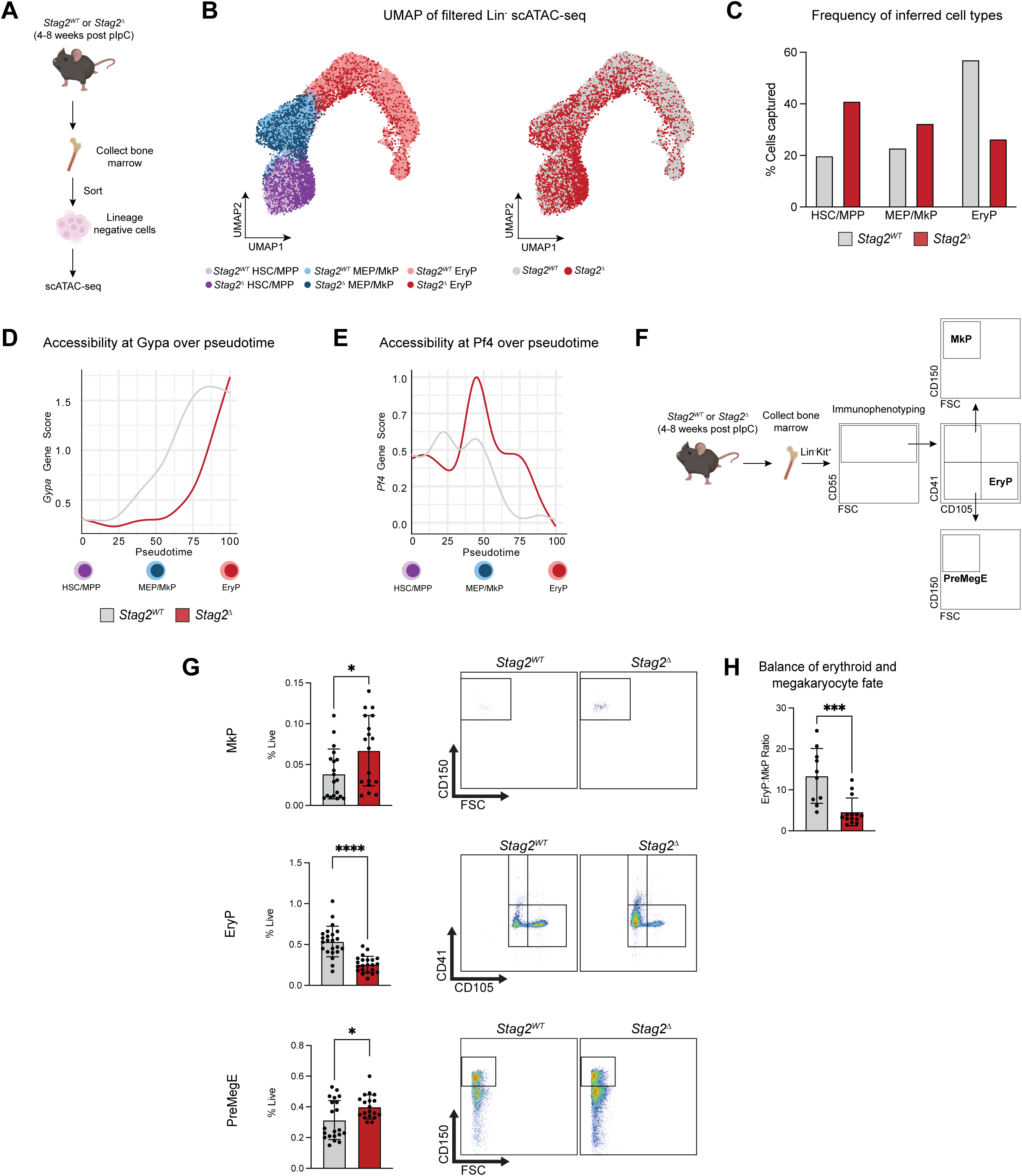
Loss of Stag2 perturbs balance of erythroid-megakaryocytic fate. (A) Schematic diagram of scATAC-seq for lineage negative hematopoietic stem and progenitor cells (B) UMAP visualization of scATAC-seq cells from sorted *Stag2^WT^* and *Stag2^Δ^* HSPCs (n = 3 for both conditions) filtered for inferred cells along erythroid differentiation with cell type annotation (left) and genotypes labelled (right). (C) Quantification of inferred cell types identified in (B). (D-E) *Gypa* (D) and *Pf4* (E) gene body accessibility over erythroid differentiation pseudotime. (F) Schematic diagram of bone marrow immunophenotyping for bipotent and unipotent erythroid and megakaryocytic progenitors. (G) Frequency and representative flow plots of MkP, EryP, and PreMegE from murine bone marrow after ACK lysis stained with antibodies against lineage markers (see supplemental table 1), Kit, CD55, CD41, CD105, and CD150. (H) Ratio of EryP to MkP. Each point represents data from an individual mouse, bar graphs depict mean ± SD. * p < 0.05, ** p < 0.01, *** p < 0.001, **** p < 0.0001 for comparison of data from *Stag2^WT^*and *Stag2^Δ^* assessed using unpaired two-tailed t test with Welch’s correction (α = 0.05) after using ROUT (Q=1%) method to remove outliers.

To determine Stag2-dependent chromatin accessibility changes during erythroid specification, we generated a pseudotime of erythroid differentiation and queried gene body accessibility at various lineage defining genes. We observed two synchronous altered states in the setting of Stag2 loss: 1) key targets of erythroid TFs with either delayed or impaired opening (**Figure 1D**) and 2) alternative TF targets that failed to restrict their accessibility as alternate lineage fate outcomes (i.e., megakaryopoiesis) are silenced (**Figure 1E**). The latter programs notably highlighted key megakaryocytic targets such as Pf4 and Fli1 (**Figure 1E, S1C)**.

### Loss of Stag2 skews MEP differentiation toward megakaryopoiesis

As prior immunophenotyping and transcriptional assessment of the stem and progenitor populations had not previously appreciated this chromatin bias towards megakaryopoiesis, we performed expanded cell surface marker flow cytometry for unipotent and bipotent megakaryocyte and erythroid progenitors, as previous described, to confirm whether this cell-intrinsic observation resulted in a population skew (**Figure 1F**)^29,30^. We observed a significant overall reduction in total MEPs consistent with prior work **(Figure S2A-B)**^22^. Stag2 loss induced a significant expansion of PreMegE and MkPs and a contraction of EryPs (**Figure 1G**). To understand the balance between unipotent erythroid and megakaryocytic fate, we calculated a ratio of EryP to MkP and observed that Stag2 loss results in a reduction in this ratio, indicating a shift away from erythroid to megakaryocytic fate (**Figure 1H**). Overall, loss of Stag2 impairs erythroid fate specification, owing to unbalanced output of lineage-specified unipotent progenitors.

### Stag2 is required for terminal erythroid differentiation and enucleation

To understand if altered lineage specification within the HSPC compartment altered terminal lineage commitment, we profiled the pools of Ter119^+^ and Kit^-^CD41^+^ cells, to capture the irreversibly erythroid and megakaryocyte committed cells **(Figure S2C)**. Consistent with our prior findings, *Stag2^Δ^* mice had an overall reduction in Ter119^+^ erythroblasts **(Figure S2D-E)**. We also observed a previously unappreciated expansion in Kit^-^CD41^+^ megakaryocytes **(Figure S2F-G)**. Given the fate imbalance at the level of the unipotent erythroid and megakaryocytic progenitors we noted, we sought to determine if the effect of Stag2 loss on chromatin accessibility was restricted to lineage specification or if chromatin accessibility was also perturbed in terminal erythroid differentiation (TED). As terminal erythropoiesis exhibits significant cell amplification, we isolated progenitor and TED populations separately to ensure sufficient representation and capture rate. In contrast to the HSPC scATAC-seq, we observed separate and parallel distribution of *Stag2^WT^* and *Stag2^Δ^* cells in Ter119^+^ scATAC-seq by UMAP dimensional reduction (**Figure 2A-B; S3A-B)**. To ascertain what differences accounted for this finding, we examined the differentially accessible genes between *Stag2^WT^* and *Stag2^Δ^* erythroblasts. We noted loss of accessibility at erythroid genes and gain of accessibility at megakaryocyte genes **(Figure S3C).** As with our HSPC scATAC-seq, we queried gene body accessibility over TED and observed two different synchronous states: 1) key erythroid differentiation genes that have impaired accessibility (**Figure 2C**) and 2) aberrant maintenance of alternative lineage (i.e., megakaryocyte) genes (**Figure 2D**). To validate this analysis in an unbiased manner, we generated terminal erythroid and megakaryocyte differentiation modules using genes known to be essential for each lineage’s ontogeny^31–33^. We observed that terminal erythroid differentiation was negatively enriched, while megakaryocyte differentiation was positively enriched amongst *Stag2^Δ^* erythroblasts (**Figure 2E**).

**Figure 2.**
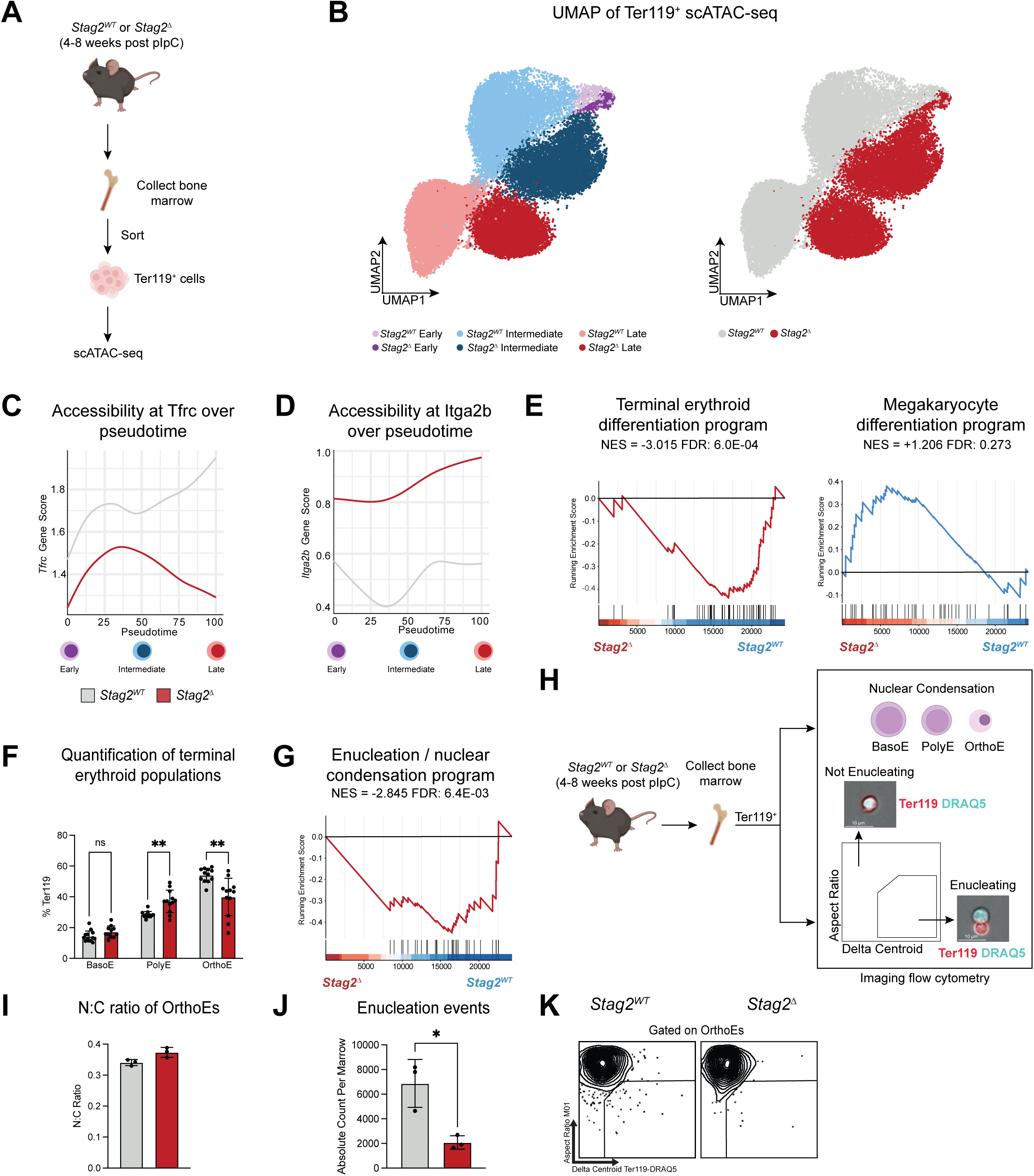
*Stag2^Δ^*impairs erythroid commitment and terminal erythroid differentiation. (A) Schematic diagram of scATAC-seq for Ter119^+^ erythroblasts (B) UMAP visualization of scATAC-seq cells from sorted *Stag2^WT^* and *Stag2^Δ^*Ter119^+^ erythroblasts (n = 3 for both conditions) with cell type annotation (left) and genotypes labelled (right). (C-D) *Tfrc* (C) and *Itga2b* (D) gene body accessibility over terminal erythroid differentiation pseudotime. (E) Gene set enrichment analysis (GSEA) plot of the terminal erythroid differentiation program (left) and megakaryocyte differentiation program (right). The plot represents the ranked, ordered, nonredundant list of genes with those on the left (red) having a higher correlation with *Stag2^Δ^*, whereas those on the right (blue) have a higher correlation with *Stag2^WT^*. The vertical black lines indicate the position of each gene in the gene set. (F) Frequency of basophilic erythroblasts (BasoE), polychromatic erythroblasts (PolyE), and orthochromatic erythroblasts (OrthoE) relative to total Ter119^+^ erythroblasts from bone marrow of *Stag2^WT^* and *Stag2^Δ^*mice after ACK lysis as determined by flow cytometry. (G) GSEA plot of the enucleation / nuclear condensation program. (H) Schematic diagram of assessing nuclear condensation and enucleation with imaging flow cytometry. (I) Quantification of median N:C ratio of OrthoE as determined by imaging flow cytometry. (J) Quantification of absolute number of enucleation events detected in bone marrow as determined by imaging flow cytometry. (K) Representative flow cytometry plot identifying actively enucleating OrthoEs stained with antibodies against Ter119, Kit, and CD71, and nuclear dyes DAPI and DRAQ5. Each point represents data from an individual mouse, bar graphs depict mean ± SD. * p < 0.05, ** p < 0.01, *** p < 0.001, **** p < 0.0001 for comparison of data from *Stag2^WT^* and *Stag2^Δ^* assessed using unpaired two-tailed t test with Welch’s correction (α = 0.05) after using ROUT (Q=1%) method to remove outliers.

Based on these observations, we hypothesized that there would be a defect in terminal erythroid differentiation associated with Stag2 loss. To test this hypothesis, we profiled each individual subpopulation within the Ter119^+^ pool to determine which specific erythroblast populations were most affected by Stag2 loss. *Stag2^Δ^* mice had significantly more polychromatic erythroblasts (PolyE) and fewer orthochromatic erythroblasts (OrthoE) (**Figure 2F).** The frequency of proerythroblast (ProE) was not significantly affected by Stag2 loss **(Figure S2H)**. As the transition from PolyE to OrthoE is hallmarked by nuclear condensation, a necessary antecedent to enucleation, we generated a module of genes previously shown to facilitate nuclear condensation and enucleation, and noted a striking negative enrichment of these genes in *Stag2^Δ^* erythroblasts (**Figure 2G**)^34^. To confirm this finding, we used imaging flow cytometry to identify these populations and interrogate nuclear condensation and enucleation dynamics in primary murine bone marrow (**Figure 2H**)^35^. We observed a significant increase in the nuclear:cytoplasmic ratio (N:C ratio) of OrthoEs (**Figure 2I**), pointing to a defect in nuclear condensation and consistent with prior reports of erythroid lineage N:C dysynchrony in histopathological assessments of *Stag2^Δ^* murine bone marrow^22^. To understand whether this increase in N:C ratio resulted in functional alteration, we quantified the number of enucleation events and observed a significant reduction in the number of enucleation events in *Stag2^Δ^* mice compared to *Stag2^WT^* (**Figure 2J-K**). Collectively, our comprehensive evaluation of the entire erythroid continuum in *Stag2^WT^* and *Stag2^Δ^*mice indicates a pan-erythropoietic requirement for Stag2.

### Stag2 loss alters chromatin accessibility and cell type-specific transcriptional fidelity in EryPs

As our chromatin and immunophenotypic data pointed towards megakaryocytic skewing associated with increased megakaryocyte-specific target gene accessibility, we next aimed to determine whether this effect was evident at unipotent erythroid commitment at the EryP cell stage. Here, megakaryocyte differentiation programs are physiologically repressed to facilitate irreversible erythroid commitment^36–38^. To determine the role of chromatin accessibility at this critical cell fate specification stage, we interrogated the accessibility and transcriptional modules in EryPs that facilitate erythroid differentiation as well as for repressing the megakaryocyte differentiation program by pseudobulking our scATAC-seq. Amongst pseudobulked EryPs, the genes that had the greatest loss of accessibility included genes such as *Gypc*, *Hba-a1*, and *Hbq1b*, while *Pawr*, *Dkk2*, *Astn2*, and *Fli1* were amongst the genes that significantly gained accessibility (**Figure 3A).** Consistent with these findings, we observe via gene set enrichment analysis (GSEA) of inferred EryPs that the erythroid differentiation program is significantly negatively enriched, while programs associated with megakaryocyte ontology are positively enriched (**Figure 3B**).

**Figure 3.**
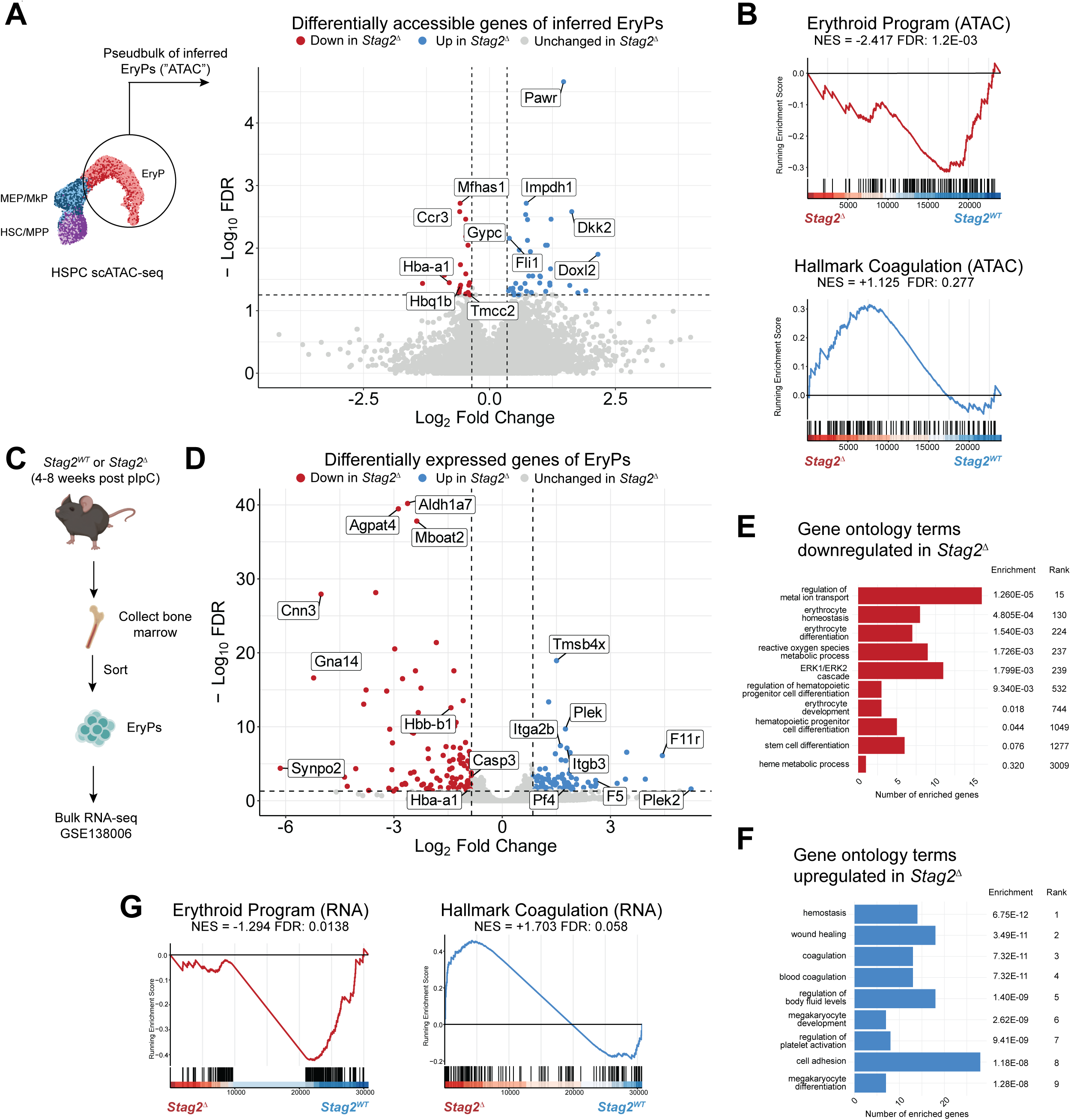
*Stag2^Δ^*results in differentially accessible and expressed erythroid and megakaryocyte programs in EryPs. (A) Volcano plot representing statistical significance (-log_10_FDR) vs. magnitude of DNA accessibility changes (log_2_FoldChange) from a comparison of inferred *Stag2^WT^* and *Stag2^Δ^* EryPs. (B) GSEA plot of the erythroid differentiation program (top) and “Hallmark Coagulation” pathway (bottom). The plot represents the ranked, ordered, nonredundant list of genes with those on the left (red) having a higher correlation with *Stag2^Δ^*, whereas those on the right (blue) have a higher correlation with *Stag2^WT^*. The vertical black lines indicate the position of each gene in the gene set. (C) Schematic diagram of bulk RNA-seq workflow. (D) Volcano plot representing statistical significance (-log_10_FDR) vs. magnitude of RNA expression changes (log_2_FoldChange) from a comparison of *Stag2^WT^*and *Stag2^Δ^* EryPs. (E-F) Bar plot of enriched gene ontology terms amongst downregulated (E) and upregulated (F) genes in *Stag2^Δ^* EryPs. (G) GSEA plot of the erythroid differentiation program (left) and “Hallmark Coagulation” pathway (right). NES, normalized enrichment score.

To assess if the noted DNA accessibility perturbations resulted in altered transcriptional output, we analyzed our already existing bulk RNA-seq of *Stag2^WT^* and *Stag2^Δ^*EryPs^22^. Indeed, we observed numerous erythroid (i.e., *Hba-a1*, *Hbb-b1*) and megakaryocytic genes (i.e., *Plek, F11r)* that are differentially expressed with loss of Stag2, concordant with our scATAC-seq findings (**Figure 3C-D**). Gene ontology (GO) analysis of differentially expressed genes revealed an overrepresentation of erythropoiesis related pathways and megakaryopoiesis related pathways within the downregulated and upregulated genes, respectively (**Figure 3E-F).** We then repeated our GSEA analyses on our RNA-seq data and observed concordant results with a significant negative enrichment in the erythroid transcriptional program and positive enrichment in the megakaryocyte transcriptional program (**Figure 3G**). Together, our pseudobulked scATAC- and RNA-seq data suggest that loss of Stag2 results in impaired induction and silencing of erythroid and megakaryocytic programs, respectively.

### Gata1 lineage specificity is disrupted by altered chromatin remodeling

To investigate the mechanism by which alterations in DNA accessibility translate into transcriptional changes, we queried the significant genes identified from our RNA-seq for TF enrichment^39^. Amongst the downregulated genes, Gata1, Maz, and CTCF were amongst the top hits (**Figure 4A**). Interestingly, Gata2, Ets, Tal1, and Gata1 were amongst the top hits within TFs enriched in upregulated genes (**Figure 4B**). Although Gata1 was identified as one of the top enriched TFs in both down- and upregulated genes, the gene targets that represented increased and decreased Gata1 engagement were distinct and non-overlapping **(Figure S4A)**. To assess whether the Gata1 targets represented unique and specific cistromes, we mapped reference Gata1 ChIP-seq experiments of erythroid and megakaryocytic cells to dichotomize genes that are bound by Gata1 in a lineage-specific manner^40^. We thus created two gene sets that are each positively regulated by Gata1 but have lineage-specific roles as either erythroid-specific Gata1 targets (“Gata1 Erythroid”) or as megakaryocyte-specific Gata1 targets (“Gata1 Megakaryocyte”) targets. Our differentially expressed genes in the *Stag2^Δ^* EryPs population were negatively enriched for Gata1 Erythroid (NES -1.85, FDR = 0.0509), while megakaryocyte-specific Gata1 targets were positively enriched (NES +2.82, FDR = 1.38E-05) (**Figure 4C**). We observed the same trend when queried against pseudobulked scATAC-seq populations (**Figure 4D**).

**Figure 4.**
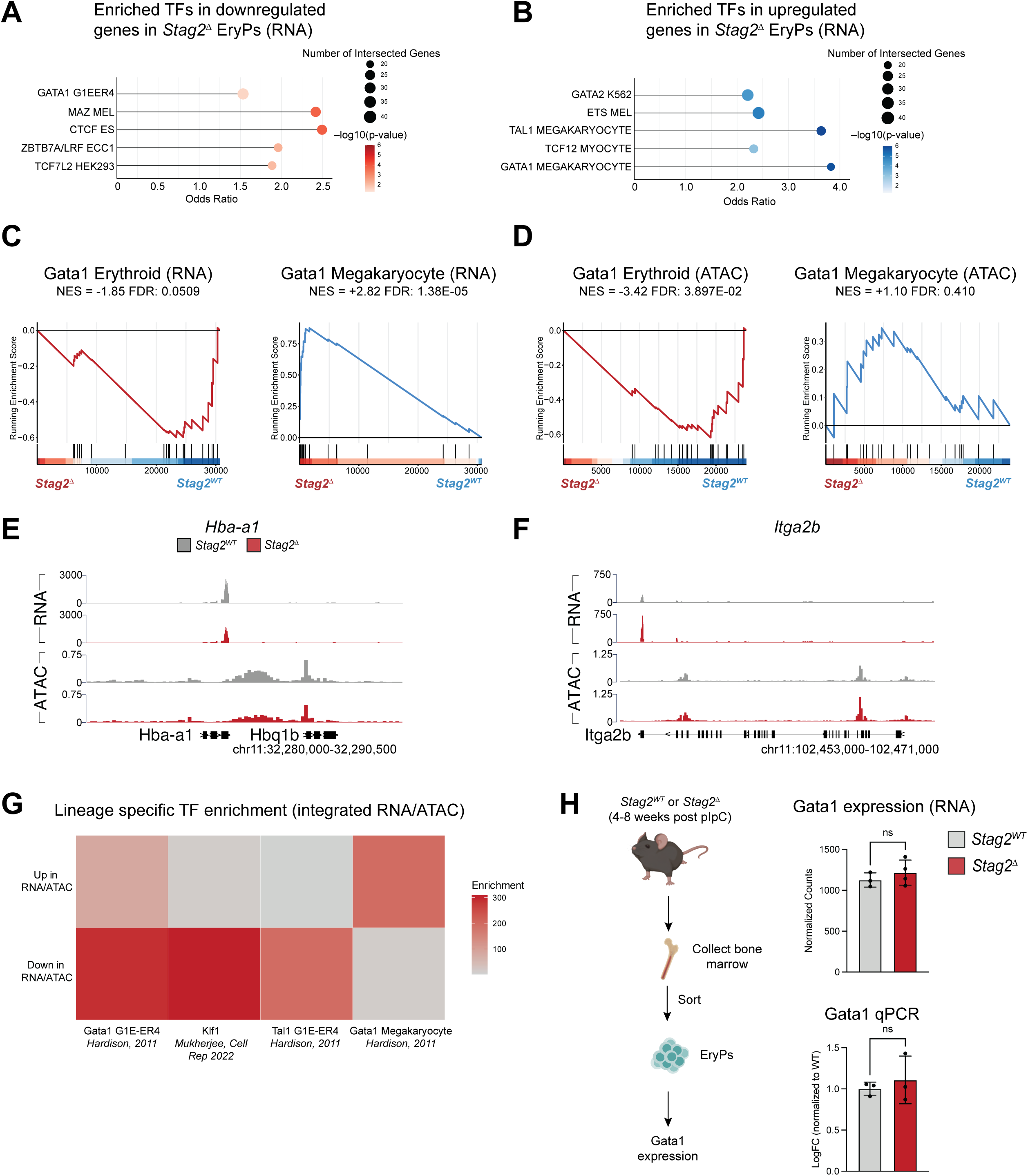
Loss of Stag2 results in altered engagement of lineage-specific Gata1 cistromes. (A-B) Lollipop plot representing transcription factor enrichment amongst downregulated (A) and upregulated (B) genes assessed via bulk RNA-seq of *Stag2^WT^*and *Stag2^Δ^* EryPs using CHEA3. (C-D) GSEA plot of the Gata1 Erythroid (left) and Gata1 Megakaryocyte (right) cistromes as determined by RNA-seq (C) and pseudobulked scATAC-seq (D). The plot represents the ranked, ordered, nonredundant list of genes with those on the left (red) having a higher correlation with *Stag2^Δ^*, whereas those on the right (blue) have a higher correlation with *Stag2^WT^*. The vertical black lines indicate the position of each gene in the gene set. (E-F) Trackplot of RNA-seq and pseudobulked scATAC-seq for *Hba-a1* (E) and *Itga2b* (F). (G) Enrichment of erythroid and megakaryocyte transcription factors amongst genes with concordant expression and accessibility changes. Transcription factor enrichment was calculated using two-tailed Fisher’s exact test (α = 0.05). (H) Schematic diagram depicting assessment of Gata1 expression levels in EryPs with bar plots representing Gata1 expression levels as determined by RNA-seq (top) and qPCR (bottom). Each point represents data from an individual mouse, bar graphs depict mean ± SD. * p < 0.05, ** p < 0.01, *** p < 0.001, **** p < 0.0001 for comparison of data from *Stag2^WT^* and *Stag2^Δ^*assessed using unpaired two-tailed t test with Welch’s correction (α = 0.05)

To more directly connect Stag2 dependent accessibility changes at Gata1 targets with transcriptomic changes, we integrated our RNA-seq with the corresponding pseudobulked scATAC-seq. Indeed, numerous genes relevant to erythroid and megakaryocytic differentiation had concordant (e.g., same direction of RNA/ATAC-seq changes) alterations in expression and accessibility (**Figure 4E-F, S4B)**. Amongst the genes that experienced concordant changes in RNA and ATAC signal, we queried TF enrichment and observed that Gata1 erythroid targets enriched to a greater degree amongst genes that lost accessibility and expression, while Gata1 megakaryocyte targets enriched to a greater degree amongst genes that gained accessibility and expression (**Figure 4G**). Furthermore, we also detected enrichment of lineage-specific TFs downstream of Gata1 with Klf1 and Tal1 enrichment amongst genes that had concordant losses (**Figure 4G**). Subsequently, we wondered if altered Gata1 stoichiometry altered cistrome engagement. To test this hypothesis, we assayed Gata1 expression levels and found no difference in Gata1 expression either in RNA-seq or by qPCR in *Stag2^Δ^* EryPs (**Figure 4H**). Collectively, these data point towards an epigenetic mechanism of altered Gata1 lineage specificity in the setting of Stag2 loss.

### Lineage-specific Gata1 cistromes are reprogrammed by chromatin state

In the absence of quantitative alterations in Gata1 expression, we hypothesized that altered chromatin accessibility results in aberrant Gata1 chromatin occupancy. To evaluate Gata1 chromatin occupancy, we conducted CUT&RUN for Gata1 in *Stag2^WT^* and *Stag2^Δ^* EryPs (**Figure 5A**). Overall, we identified 80,721 Gata1 bound peaks in *Stag2^WT^* EryPs and 54,251 peaks in *Stag2^Δ^* EryPs (**Figure 5B**). Next, we identified genes with significantly different Gata1 occupancy with Stag2 loss. Among the top genes that gained Gata1 occupancy included *Mx1*, *Dip2c,* and *Rhob*, while top genes that lost Gata1 occupancy included *Pdgfd*, *Noct*, *Tpk1*, and *Ebp42* (**Figure 5C-E**). GO analysis of genes that lost Gata1 occupancy revealed overrepresentation of erythroid related pathways and megakaryocyte related pathways amongst genes that gained Gata1 occupancy, consistent with our RNA- and pseudobulked scATAC-seq data (**Figure 5F-G**). Together, these analyses provide evidence that loss of Stag2 alters EryP Gata1 target occupancy, which we hypothesized may lead to ectopic maintenance of accessible megakaryocyte targets.

**Figure 5.**
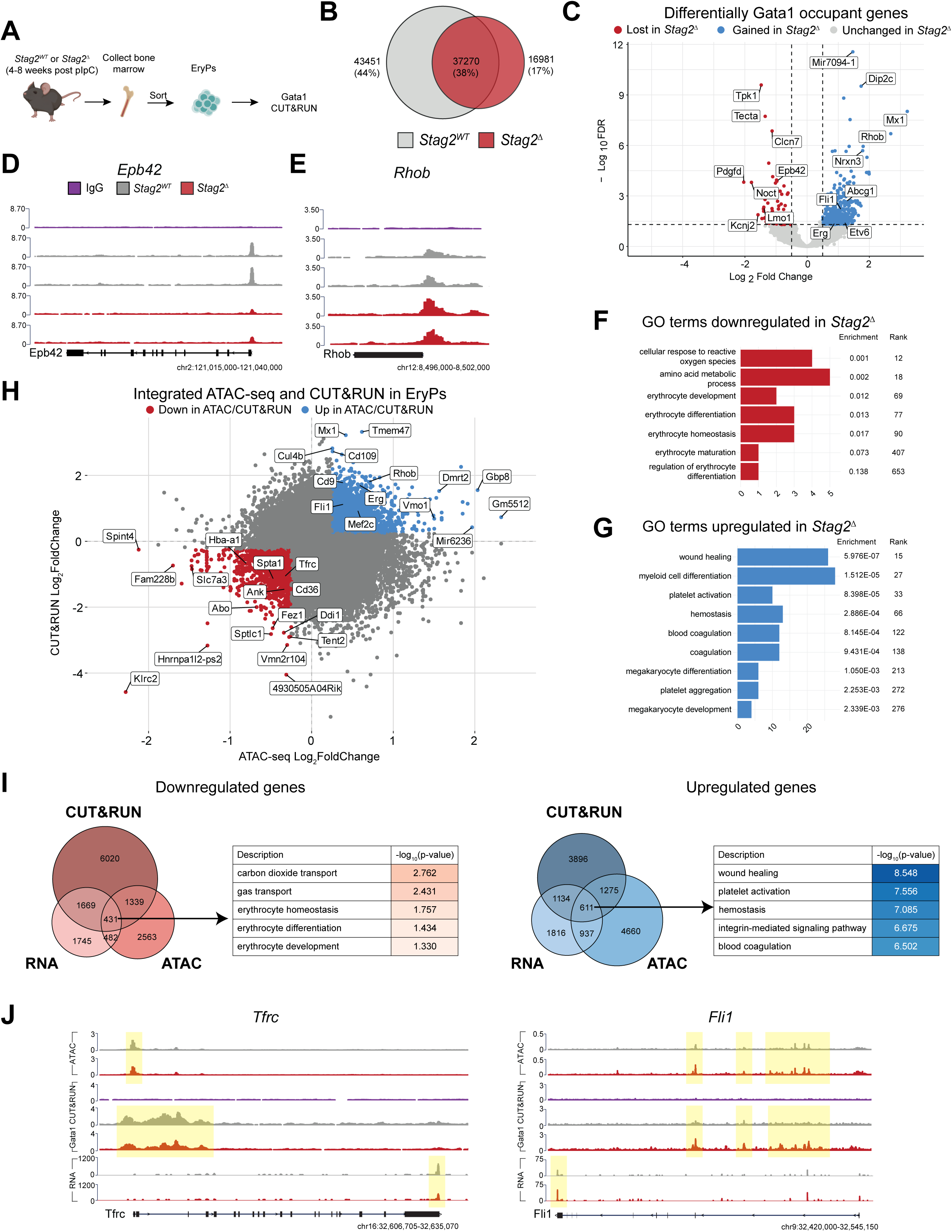
*Stag2^Δ^*alters Gata1 occupancy at lineage-specific targets. (A) Schematic diagram of Gata1 CUT&RUN workflow. (B) Venn diagram depicting the number of Gata1 bound peaks called in *Stag2^WT^* and *Stag2^Δ^* EryPs. (C) Volcano plot representing statistical significance (-log_10_FDR) vs. magnitude of Gata1 occupancy changes (log_2_FoldChange) from a comparison of *Stag2^WT^*and *Stag2^Δ^* EryPs. Only the largest peak per gene is labelled. (D-E) Trackplots displaying Gata1 occupancy at *Epb42* (F) and *Rhob* (G). (F-G) Bar plot of enriched gene ontology terms amongst genes losing (F) and gaining (G) Gata1 occupancy in *Stag2^Δ^*EryPs. (H) Plot of magnitude of Gata1 occupancy changes vs. magnitude of DNA accessibility changes in *Stag2^WT^* and *Stag2^Δ^* EryPs. (I) Venn diagram of overlap of RNA expression, DNA accessibility, and Gata1 occupancy changes per gene. Table to the right represents gene ontology terms enriched among the genes that experience concordant expression, accessibility, and occupancy changes. (J) Trackplots displaying DNA accessibility, Gata1 occupancy, and RNA expression at *Fli1* and *Tfrc*.

To determine whether differential accessibility functionally results in non-erythroid Gata1 target engagement, we integrated our Gata1 CUT&RUN data with our pseudobulked scATAC-seq. Numerous erythroid (i.e., *Spta1*, *Tfrc*, *Ank*, *Cd36*, *Abo*, *Hba-a1)* genes were among those that lost accessibility and Gata1 occupancy and numerous megakaryocyte genes (i.e., *Rhob, Cd109, Cd9, Fli1, Erg*) were among those that gained accessibility and Gata1 occupancy (**Figure 5H**). To identify a core set of genes that had concordant RNA, ATAC, and Gata1 occupancy changes, we integrated all three of our sequencing datasets. We identified 431 genes (i.e., *Tfrc*, *Hba-a1*) that lost and 611 genes (i.e., *Fli1*, *Cd9*) that gained accessibility, Gata1 occupancy, and expression (**Figure 5I-J**). Gene ontology of the genes at the intersection of all three sequencing techniques identified erythroid and megakaryocyte pathways amongst the downregulated and upregulated gene sets, respectively (**Figure 5I**). Collectively, these data highlight that loss of Stag2 in EryPs confers an altered chromatin conformation that fails to restrict Gata1 chromatin binding to erythroid-specific targets and enables ectopic megakaryocyte-specific Gata1 transcriptional programs to impair erythropoiesis.

### Stag2-deficient erythroid progenitors exhibit impaired erythropoiesis and ectopic megakaryopoiesis

Finally, to understand the functional implications of discordant DNA accessibility and alteration of the erythroid Gata1 cistrome on lineage fidelity and differentiation capacity, we sorted *Stag2^WT^* and *Stag2^Δ^*EryPs and seeded them into liquid media supplemented with SCF (100 ng/mL) and EPO (4U/mL; **Figure 6A**). After 48 hours, we observed fewer cells overall **(Figure S5A-C)** and significantly fewer Ter119^+^CD71^+^ cells generated by *Stag2^Δ^*EryPs compared to *Stag2^WT^* EryPs (p < 0.05; **Figure 6B-C**). Additionally, these *Stag2^Δ^* EryPs produced significantly more CD41^+^CD42d^+^ megakaryocytes (p < 0.05; **Figure 6D**). To confirm these findings in an orthogonal assay, we assessed colony forming capacity of *Stag2^WT^* and *Stag2^Δ^* EryPs in either erythroid directing methylcellulose or pan-myeloid directing methylcellulose (**Figure 6E**). In erythroid-directing methylcellulose, we observed significantly fewer erythroid colonies generated by *Stag2^Δ^* EryPs compared to *Stag2^WT^*EryPs (p < 0.05; **Figure 6F-H**). In pan-myeloid methylcellulose, we observed a significant reduction in Ter119^+^CD71^+^ cells and an increase in CD41^+^ cells (p < 0.05; **Figure 6I-K**). Collectively, these data provide a link between the transcriptional, accessibility, and Gata1 occupancy aberrancies to impaired EryP function in the context of Stag2 loss.

**Figure 6.**
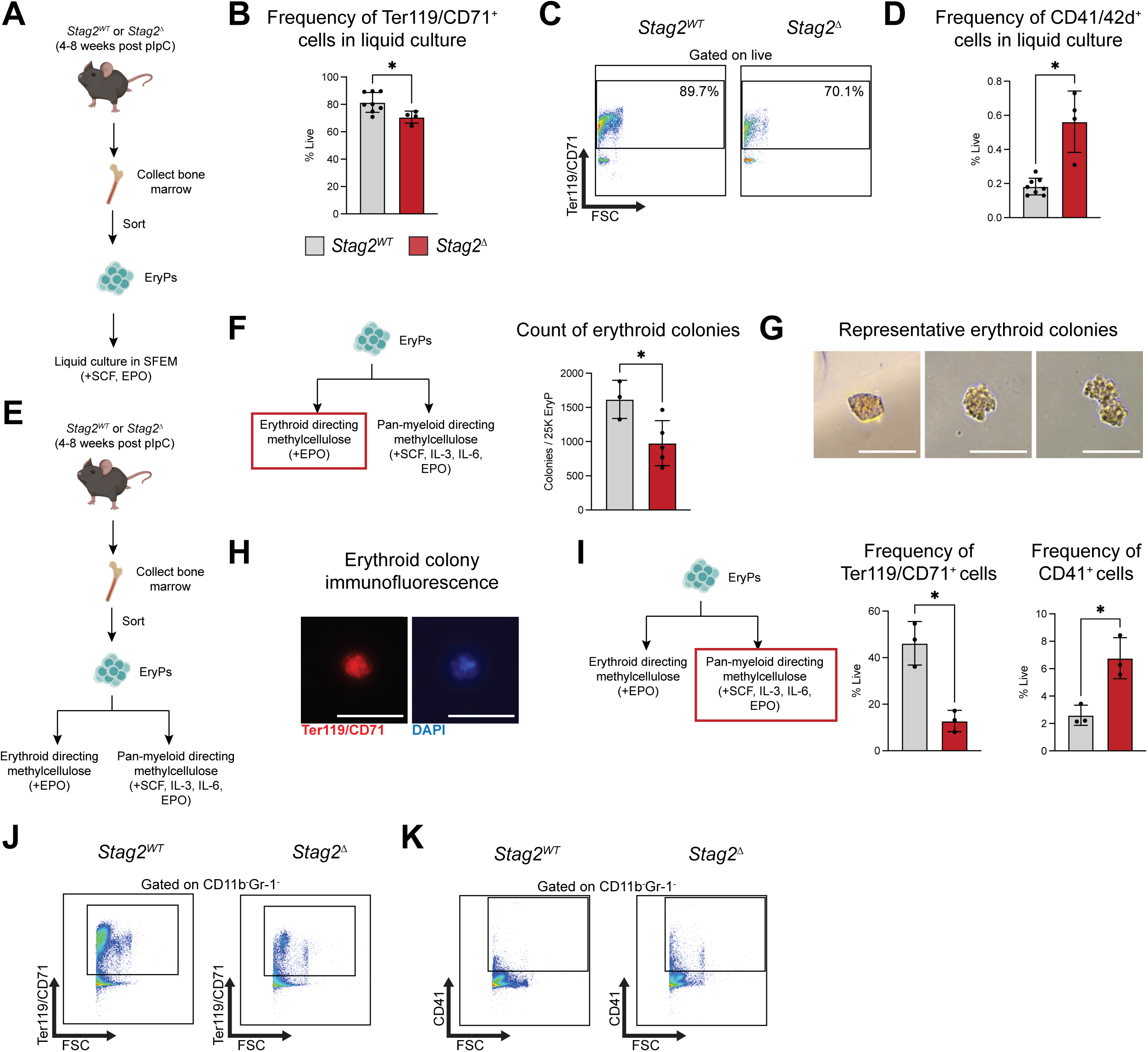
Stag2-deficient EryPs have diminished erythroid output and increased megakaryocyte output. (A) Graphical depiction of liquid culture differentiation workflow. (B) Frequency of Ter119^+^CD71^+^ cells generated by EryPs after 48 hours in liquid culture as determined by flow cytometry. (C) Representative flow cytometry plots identifying Ter119^+^CD71^+^ cells stained with antibodies against Ter119, CD71, CD41, CD42d, and Kit. (D) Frequency of CD41^+^42d^+^ cells generated by EryPs after 48 hours in liquid culture as determined by flow cytometry. (E) Graphical depiction of methylcellulose assay workflow. (F) Absolute count of the number of erythroid colonies generated by EryPs after 48 hours in erythroid directing methylcellulose identified by brightfield microscopy. (G) Representative brightfield microscopy images of erythroid colonies identified. Images were obtained using a 40X/0.55 achromatic objective lens. The scale bar represents 50 µm. (H) Representative immunofluorescence images of erythroid colonies generated by EryPs after 48 hours in erythroid directing methylcellulose. Images were obtained using a 20X/0.45 objective lens. The scale bar represents 50 µm. (I) Frequency of Ter119^+^CD71^+^ (left) and CD41^+^ (right) cells generated by EryPs after 8 days in pan-myeloid directing methylcellulose as determined by flow cytometry. (J-K) Representative flow cytometry plots identifying Ter119^+^CD71^+^ (J) and CD41^+^ (K) cells stained with antibodies against CD11b, Gr-1, CD41, Ter119, and CD71. Each point represents data from an individual mouse, bar graphs depict mean ± SD. * p < 0.05, ** p < 0.01, *** p < 0.001, **** p < 0.0001 for comparison of data from *Stag2^WT^* and *Stag2^Δ^* assessed using unpaired two-tailed t test with Welch’s correction (α = 0.05) after using ROUT (Q=1%) method to remove outliers.

### STAG2-deficient HSPCs undergo dysplastic erythropoiesis and recapitulate essential features of murine model

As STAG2 is a recurrent mutational target in MDS, we sought to determine the generalizability of our murine observations of Stag2 mutant dyserythropoiesis in human cells and MDS patients. Using human CD34^+^ HSPCs isolated from umbilical cord blood, we conducted a shRNA knockdown of STAG2 (**Figure 7A-B**). Flow cytometric analyses identified delayed upregulation of CD36 and gain of CD235, indicating delayed erythroid specification and terminal commitment (**Figure 7C-D**)^27,41^. We then assessed terminal erythroid differentiation. Here, STAG2-deficient HSPCs progressed through terminal erythroid differentiation less efficiently and had reduced enucleation (**Figure 7E-F**). To determine if the features of erythroid dysplasia were conserved in human cells, we conducted cytospins of differentiating STAG2-deficient HSPCs and observed features of myelodysplasia such as multinucleated erythroblasts and red blood cells, unique pathological morphologies described in MDS (**Figure 7G**).

**Figure 7.**
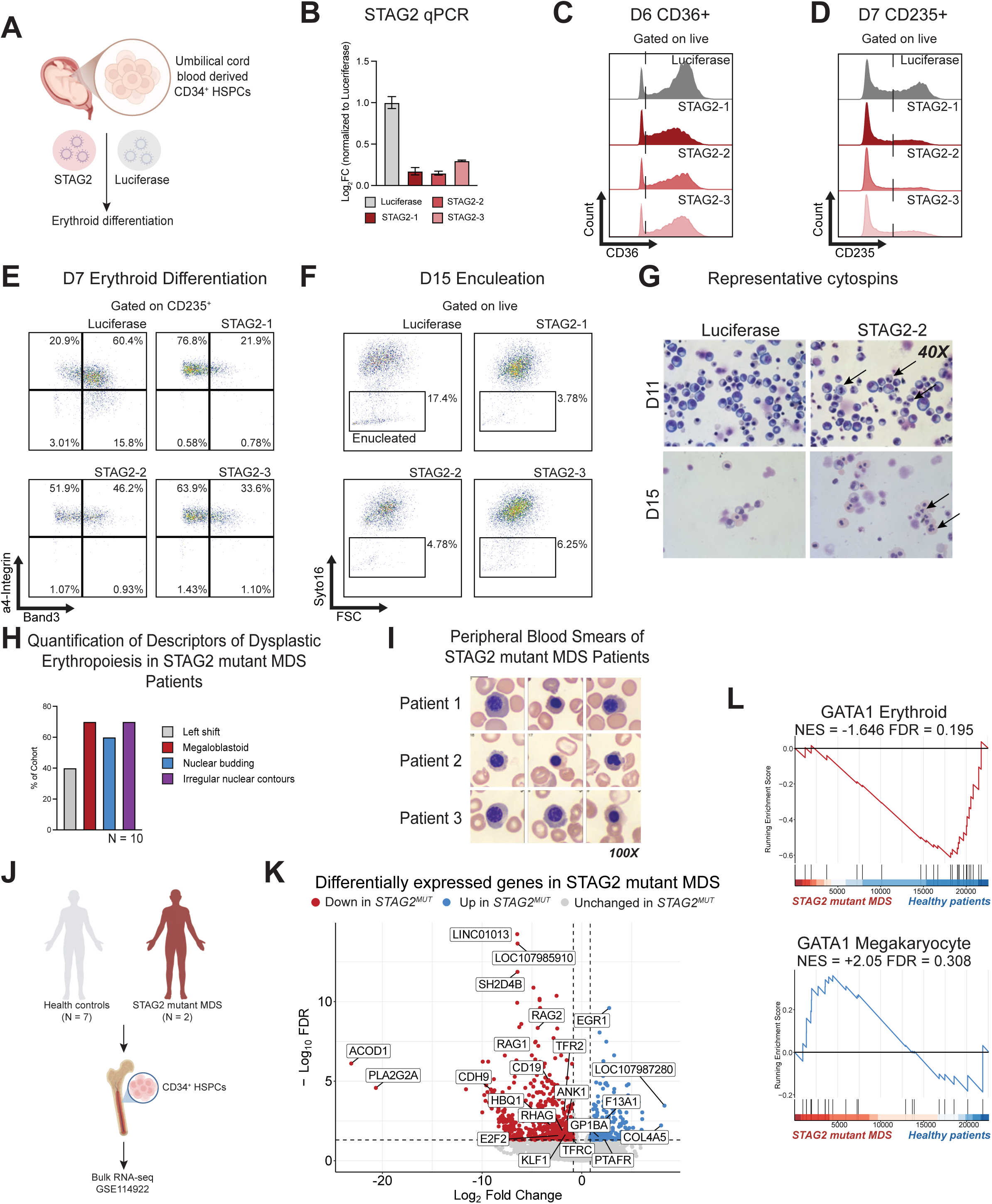
Human models of erythropoiesis and STAG2 mutant MDS patients share phenotypic and molecular features with *Stag2^Δ^* mice. (A) Schematic diagram of umbilical cord blood derived CD34^+^ HSPC shRNA knockdown and erythroid differentiation. (B) Quantification of STAG2 mRNA levels after shRNA delivery as determined by qPCR. (C) Representative flow cytometry histogram of CD36 cell surface expression of differentiating HSPCs on day 6 of erythroid differentiation. (D) Representative flow cytometry histogram of CD235 cell surface expression of differentiating HSPCs on day 7 of erythroid differentiation. (E) Representative flow cytometry plots of a4-integrin and band3 expression on D7 of in-vitro differentiation. (F) Representative flow cytometry plots of Syto16 positivity on D15 of in-vitro differentiation. (G) Representative cytospins of shLuciferase and shSTAG2 cells on D11 and D15 of in-vitro differentiation. Arrows indicate dysplastic cells. Original magnification x 40. (H) Percentage of STAG2 mutant MDS patients (N = 10) with descriptors of dysplastic erythropoiesis. (I) Representative peripheral blood smears from STAG2 mutant MDS patients obtained from CellaVision. Original magnification x 100. (J) Schematic diagram of STAG2 mutant MDS patient bulk RNA-seq. (K) Volcano plot representing statistical significance (-log_10_FDR) vs. magnitude of RNA expression changes (log_2_FoldChange) from a comparison of *STAG2^WT^*and *STAG2^MUT^* patients. (L) GSEA plot of the GATA1 Erythroid (top) and GATA2 Megakaryocyte (bottom) cistromes as determined by RNA-seq. The plot represents the ranked, ordered, nonredundant list of genes with those on the left (red) having a higher correlation with *STAG2^MUT^* patients, whereas those on the right (blue) have a higher correlation with *STAG2^WT^* patients. The vertical black lines indicate the position of each gene in the gene set.

Leveraging our annotated MDS patient cohort, we retrospectively queried our STAG2 mutant MDS patient cohort to determine if STAG2 mutant MDS patients also had features of abnormal nuclear condensation. We examined bone marrow hematopathology of MDS patients with STAG2 mutations (N = 10) and found that the majority of these patients had altered nuclear morphology (**Figure 7H**), and many had nucleated red blood cells and dysplastic erythroblasts in peripheral blood (**Figure 7I**).

Finally, to ascertain if features of altered GATA1 lineage specificity were present in primary MDS patients, we queried public RNA-seq datasets obtained from STAG2 mutant MDS patients (**Figure 7J**)^42^. Consistent with our findings, we observed that STAG2 mutant MDS patients had reduced expression of erythroid genes, increased expression of megakaryocyte genes, and dysregulation of lineage-specific GATA1 cistromes (**Figure 7K-L).** Collectively, these data support that dyserythropoiesis is a common hallmark of Stag2 loss in human models of erythropoiesis and MDS patients with shared phenotypic and molecular features of impaired erythropoiesis.

## Discussion

Our study reveals a critical role for Stag2-mediated chromatin remodeling in enforcing the erythroid-specific Gata1 cistrome and maintaining balanced erythro-megakaryopoiesis. By integrating chromatin accessibility, transcriptional profiling, and Gata1 occupancy data, we demonstrate that Stag2 loss disrupts the chromatin landscape in erythroid progenitors (EryPs), leading to a misallocation of Gata1 binding and a shift in lineage output. Our present study enabled novel identification of Gata1 as a major transcription factor (TF) affected by Stag2 loss in an *in vivo* murine model of myelodysplastic syndrome (MDS)^22,43^. Our findings specifically highlight the regulatory function of chromatin accessibility in directing TF cistrome specificity. The present findings further highlight how disordered chromatin can act as a functional driver of epigenetic dysregulation leading to imprecise lineage commitment in cell type-specific hematopoietic differentiation.

A central insight from our work is that Gata1, despite being expressed across multiple hematopoietic lineages, is highly influenced by cell type-specific chromatin structure to execute lineage-specific transcriptional programs. This positions chromatin accessibility as an active and non-redundant regulatory mechanism instructing and guiding TF binding specificity and function. These data also provide a conceptual framework for how TFs with broad expression patterns achieve distinct cistromic capabilities that likely intersect with co-binding patterns and expression levels. Notably, altered TF engagement is appreciated as a driver of human disease such as in Down Syndrome associated AML (DS-AML) where the GATA1s mutation results in preferential occupancy at canonical megakaryocyte GATA1 targets^44^. In fact, STAG2 mutations occur in 50% of patients with DS-AML, highlighting that there may be some shared oncogenic mechanism that remains to elucidated^21^. Our future studies will aim to expressly understand the cooperativity between STAG2 and GATA1s, as these findings share conspicuous immunophenotypic and epigenetic similarities to Gata1s mice such as impaired erythropoiesis and augmented megakaryopoiesis^44,45^. Consistent with this, RNA-seq of these mice revealed a negative enrichment of erythroid pathways and positive enrichment of megakaryocyte/platelet pathways^44^. Subsequent ATAC-seq and CUT&RUN revealed that Gata1s resulted in gained accessibility and chromatin occupancy at megakaryocyte genes, while erythroid genes experienced losses. Together, these studies underscore the concept of a “TF-cistrome to chromatin mismatch” as a key driver of disordered hematopoiesis. While additional work in the context of MDS and DS-AML is needed to understand the cooperativity of STAG2 mutations with other somatic mutations, the mismatch between TF availability and chromatin state may represent a generalizable mechanism by which mutations in chromatin regulators contribute to hematologic disease.

In the context of MDS, dysregulation of GATA1 kinetics and splicing has been shown to contribute to the dyserythropoiesis that is archetypal of MDS^46,47^. In fact, preliminary data from other groups has implicated impaired GATA1/KLF1 induction as a feature of MDS and one that is transiently ameliorated by drugs like luspatercept^48^. Furthermore, this may provide a reasonable hypothesis as to the transient nature of the transfusion independence obtained with luspatercept as it does not directly correct MDS associated chromatin architectural abnormalities. Hence, therapeutic strategies aimed at restoring chromatin balance or correcting TF mislocalization could help re-establish lineage fidelity and correct ineffective erythropoiesis, which is a major driver of mortality and morbidity in patients with MDS^49^.

As we, and others, establish key mechanisms driving the tumorigenic potential of STAG2 loss, a common observation is strong dysregulation of chromatin accessibility, with more muted effects on DNA looping, transcriptional control, and sister chromatid cohesion – the canonical functions of cohesin. These data demonstrate the downstream mechanistic phenomenology of altered chromatin accessibility, yet future studies are needed to understand STAG2’s noncanonical control of cell type-specific DNA accessibility. Drawing from insights in immortalized cell line models investigating other epigenetic machinery, we posit that STAG2 likely functions in concert with other epigenetic complexes such as SWI/SNF and NuRD, though its unique interactions with these complexes (in contrast to other cohesin subunits) and exact position in the cascade of epigenetic events occurring during hematopoiesis is unclear^50–52^.

In summary, we identify Stag2 as a key enforcer of erythroid identity through its role in shaping the chromatin landscape that guides Gata1 binding. Our findings underscore the necessity of coordinated TF and chromatin interactions in lineage specification and highlight TF-cistrome/chromatin mismatch as a novel paradigm in the pathogenesis of hematopoietic disorders.

## Acknowledgements

We are grateful to members of the Viny Lab, Columbia Stem Cell Initiative (CSCI), and CSCI Flow Cytometry Core for their discussion of this work and advisory technical support. This work was funded by NIH/NCI grant R37CA28657 (A.D.V.), Edward P. Evans MDS Foundation Discovery Research Grant (A.D.V.), and American Society of Hematology (ASH) Scholar Award (A.D.V).

This research was funded in part through the NIH/NCI Cancer Center Support Grant (P30CA012696), supported by the NCATS Clinical and Translational Science Awards (CTSA) grant (UL1TR001873), and used the Genomics and High Throughput Screening Shared Resource. Conventional and imaging flow cytometry experiments were performed on instruments supported by the NIH grant S10OD026845 and the Milky Way Foundation. V.S.S. is an American Society of Clinical Investigation (ASCI) PSSF Fellow and an American Society of Hematology (ASH) HONORS recipient. V.S. is a recipient of a postdoctoral fellowship from the American Italian Cancer Foundation. J.J.X. is supported by a postdoctoral fellowship from the Walker Family and the Edward P. Evans Columbia MDS Center. A.D.V is a Damon-Runyon-Doris Duke Clinical Investigator supported by the Damon Runyon Cancer Research Foundation (CI-120-22) and a Clinician Scientist Development Award from the Doris Duke Charitable Foundation.

Schematics included in the figures were generated using BioRender.

## Author’s Contributions

**V.S.S:** conceptualization, data curation, formal analysis, funding acquisition, investigation, methodology, project administration, software, validation, visualization, writing-original draft, writing-review and editing **E.M.S:** conceptualization, data curation, formal analysis, investigation, methodology, software, validation, visualization, writing-original draft, writing-review and editing **Y.C:** data curation, formal analysis, investigation, software, validation, visualization, writing-original draft **H.Y:** investigation, methodology **S.F:** investigation, resources **V.S:** investigation, supervision, validation, writing-review and editing **J.J.X:** conceptualization, investigation, software, supervision, writing-review and editing **M.N:** supervision, writing-review and editing **A.D.V:** conceptualization, data curation, funding acquisition, methodology, project administration, resources, software, supervision, visualization, writing-original draft , writing-review and editing.

