## Supplemental Methods and Figures for "Stag2 dependent chromatin remodeling enforces the erythroid-specific Gata1 cistrome"

##### *Flow cytometry and cell sorting*

For immunophenotyping, bone marrow (BM) cells were obtained by crushing femurs, tibias, and pelvises in flow cytometry buffer (FACS buffer) composed of phosphate-buffered saline (PBS) and 2% heat-inactivated fetal bovine serum (FBS, #MT35016CV, Corning). BM RBCs were removed by lysis with ACK (150 mM $NH_4Cl$, 10 mM $KHCO_3$, and 0.1 mM $Na_2EDTA$) and washed once with FACS buffer. For cell sorting, BM cells were obtained as described above and were enriched for HSPCs (lineage negative) via immunomagnetic separation (Easy™ Mouse Hematopoietic Progenitor Cell Isolation Kit, StemCell). For colony forming assays, colonies were incubated in 2 mL of warm PBS for 10 minutes and washed once with FACS buffer. Subsequently, unfractionated or HSPC enriched BM cells or unfractionated cells from in vitro liquid culture or colony forming assays were incubated with the appropriate antibody panel (see supplemental tables 2-5 for detailed panels and dilutions) for 30 minutes. Before data acquisition, stained cells were washed once with FACS buffer and resuspended in FACS buffer containing propidium iodide (BioLegend, 421301), 4',6-amidino-2-phenylindole (DAPI; Invitrogen, D1306), or 7-aminoactinomycin D (7-AAD; Invitrogen, A1310) for dead cell exclusion. Immunophenotyping data was acquired on Novocyte Penteon, Sony MA900, or Becton Dickinson (BD) FACSCanto, while cell sorting was performed on BD FACS ARIA II SORP, Sony MA900, or BD MOFLO. Data was exported in the .fcs file format and analyzed using FlowJo (v10).

##### *Imaging flow cytometry*

BM cells were isolated, and BM RBCs were lysed as described above. Subsequently, $10^7$ BM cells were stained with the appropriate antibody panel (see supplemental table 6 for detailed panels and dilutions) for 30 minutes. Stained cells were washed once with FACS buffer and resuspended in FACS buffer containing DAPI (1:1000) and DRAQ5 (eBioscience, 65-0880-92, 1:2000) for dead cell exclusion and nuclear staining, respectively as previously described[1]. Data was acquired on Cytek Amnis ImageStream X Mk II Imaging Flow Cytometer at 40X magnification using the Amnis INSPIRE software. Acquired data was then processed and analyzed using Amnis IDEAS.

##### *Immunofluorescence*

For colony forming assays, fluorophore conjugated antibodies (see supplemental table 7 for detailed panels and dilutions) were diluted in 300 µL Iscove's Modified Dulbecco's Medium (IMDM) supplemented with 1% penicillin/streptomycin, added to each well, and incubated at 37˚C at 5% $CO_2$ overnight. Data was acquired using a Keyence BZ-X810 Fluorescent Microscope at 20X original magnification.

##### *Human CD34$^+$ HSPC liquid culture and shRNA knockdown of STAG2*

CD34$^+$ cells were isolated from umbilical cord blood and cultured as previously described[2,3]. Briefly, a three phase cell culture system was used with a base medium comprised of IMDM, 2% human peripheral blood plasma (StemCell Technologies, #70039.5), 3% human type AB serum (Atlanta Biologicals, #S40110H), holo-transferrin (200 µg/mL, Sigma-Aldrich, #T4132-1G), heparin (3 IU/mL, StemCell Technologies, #07980), and insulin (10 µg/mL, Sigma-Aldrich, #I9278), supplemented with recombinant human SCF (10 ng/mL, StemCell Technologies, #78062), recombinant human IL-3 (1 ng/mL, StemCell Technologies #78040), and recombinant human erythropoietin (3 U/mL, ThermoFisher Scientific, #PHC2054).

The sequences of STAG2 shRNAs are provided in supplemental table 8**.** Briefly, 500,000 CD34$^+$ cells were used for transduction with 30 x $10^6$ transduction units of virus[3]. Cells were transduced by centrifuging at 1500 g at room temperature for 2 hours, incubated overnight at 37˚C in a humidified incubator, washed, and resuspended in fresh medium. After 24 hours of culture, puromycin (1 µg/mL) was added to culture until D11 of culture.

*Cytospin preparation*
Cytospins from human CD34$^+$ HSPC liquid culture were prepared and imaged as previously described[3]. Briefly, 100,000 cells were used to prepare cytospins, successively stained with May-Grunwald (Sigma MG500) and Giemsa (Sigma GS500) stains and imaged using a Leica DM2000 inverted microscope.

*Quantitative real-time PCR* (qPCR)
For mouse EryPs, total RNA was isolated from 35,000 cells using the RNeasy mini kit (Qiagen). RNA concentrations were determined using Qubit fluorometric quantification (ThermoFisher) and reverse transcribed to cDNA using the Verso cDNA Synthesis kit (ThermoFisher). qPCR was performed using TaqMan Universal PCR master mix master mix (Applied Biosystems) and TaqMan targeting Gata1 (Mm01352636_m1) on CFX Opus Real-Time PCR systems (BioRad). Gata1 expression levels were normalized to Gapdh (Mm99999915_g1).

For human CD34$^+$ HSPCs, total RNA was isolated and quantified as described above. Reverse transcription and qPCR were performed as previously described[3].

*RNA sequencing data analysis*
For bulk RNA-seq, the normalized count matrix was downloaded from GEO and input to DESeq2 (1.46.0). Normalized expression values were estimated using transcripts per million (TPM). Significantly differentially expressed genes were identified based on a FDR threshold < 0.05 and a $Log_2$FoldChange (LFC) cutoff of 0.85. Gene ontology analysis and gene set enrichment analysis were performed using clusterProfiler (v.14.6) using gene sets from the Molecular Signatures Database (category M2)[4,5]. Custom defined gene sets were curated by using already existing bulk RNA-seq data[22] (erythroid program corresponds to the top 200 genes upregulated during the transition from LSK to EryP), studies of erythroid and megakaryocyte ontology (terminal erythroid differentiation program, megakaryocyte differentiation program, enucleation / nuclear condensation program), or lineage specific ChIP-seq datasets (lineage specific Gata1 cistromes). These gene sets are provided in supplemental table 9. Transcription factor enrichment was performed using CHEA3[6]. Enriched transcription factors with at least 15 intersecting targets and the lowest FDR values were then ranked by the number of intersecting targets.

*scATAC sequencing and data analysis*

Cells were pelleted at 300 g at 4˚C for 5 minutes and resuspended in PBS supplemented with 0.04% BSA. Cells were lysed, washed, and nuclei were isolated as per manufacturer's instructions (10X Genomics). Isolated nuclei were supplied to the JP Sulzberger Columbia Genome Center Single Cell Core and sequenced on a NovaSeq 6000 to a depth of 250-300M reads (HSPC) or NovaSeq X to a depth of 380-435M reads (TED).

Raw sequencing reads were aligned to the mm10 reference genome and pre-processed by the Columbia Genome Center Single Cell Core using Cell Ranger ARC (2.0.2). Arrow files were generated from the processed scATAC-seq fragments using ArchR (1.0.3)[7]. Cells with fewer than 1000 fragments and TSS < 4 were filtered out, resulting in 87,366 cells with a median number of 7567 fragments across 6 samples (TED) and 18,840 cells with a median number of 8142 fragments across 6 samples (HSPC). Automatic detection of doublets using ArchR's *addDoubletScores* was used with default parameters, removing 10-15% of cells per sample. Dimensionality reduction was performed using Iterative Latent Semantic Indexing (LSI), an approach adapted from natural language processing which is ideal for sparse and noisy data such as scATAC-seq data. ArchR's approach also performs a depth normalization per single cell, ensuring that features are weighed based on their relative frequency. To visualize the data, a UMAP projection was generated from the first 30 LSI components, using 30-nearest-neighbor graphs (cosine similarity) with a minimum distance parameter of 0.5.

To determine appropriately defined clusters, an iterative process of Leiden clustering and cell type annotation was conducted to guide cluster identification towards biologically meaningful definitions. Narrow cluster definitions were initially made with a high resolution of 3, and carefully, the resolution parameter was loosened until visualization of marker gene scores indicated clear demarcation between clusters (a final resolution of 0.8). Gene scoring controlled for TSS enrichment and log-transformed unique fragment counts as covariates, pseudobulking by the latest Leiden clustering. All non-erythrocyte lineage cell types were removed from the analysis, and the graphs were regenerated using the updated dataset.

Peaks in each annotated cell type (per genotype) were combined to form distinct group coverages, which were then used for differential gene and module scoring. Differential accessibility was assessed using the Wilcoxon signed-rank test, adjusting for covariates (TSS enrichment, log-transformed unique fragment counts) as done above. Significantly differentially accessible genes were identified based on a FDR threshold < 0.05 and a $Log_2$FoldChange cutoff of 0.35. Clusters containing cells of interest were then pseudobulked using ArchR, exported, and GSEA was performed in R as described above.

To compare the trajectories of each genotype, the ArchR function *plotTrajectory* was extended to visualize two distinct trajectories on the same plot. The psuedotimes were defined, smoothed and graphed in a similar manner to the original function - using *addTrajectory* for each genotype individually and a predefined trajectory backbone of the lineage cell types as a rough ordering of cells. From this, ArchR fits a smoothed continuous trajectory, scaled to 100, such that both trajectories can coexist meaningfully on the same plot.

##### *GATA1 CUT&RUN and data analysis*

CUT&RUN was conducted using 500,000 EryPs per target, rabbit anti-GATA1 antibody (Abcam, ab11852, 1.5 µg/sample), and 5 ng of CUT&RUN DNA for library preparation. Library quantity and size distribution were verified on a D5000 tape using an Agilent 4150 Tapestation. CUT&RUN libraries were sequenced on a NovaSeq X to a depth of 30M reads.

Sequencing reads were processed with Nextflow using the nf-core/cutandrun pipeline (v3.2.2; nf-core/tools v2.12)[8,9]. Raw FASTQ replicates were concatenated and subjected to QC with FastQC (v12.1) and aggregated in MultiQC (v1.17). Then adapter and quality trimming (Q ≥ 20, length ≥ 20 nt) was performed with Trim Galore! (v0.6.7; Cutadapt v2.10). Reads were aligned to mm10 with Bowtie 2 (v2.4.5), filtered for MAPQ ≥ 30, multimappers, mitochondrial reads and ENCODE blacklist regions with samtools (v1.16.1), and PCR duplicates marked via Picard (v3.1.0)[10]. BedGraph coverage was generated with bedtools (v2.30.0) and converted to bigWig using bedGraphToBigWig (UCSC v4)[11]. Narrow peaks were called with MACS2 (v2.2.7.1) and SEACR (v1.3) respectively[12,13]. Library complexity was estimated with preseq (v2.0.3), fragment-size distributions, FRiP scores, promoter-proximal enrichment, and replicate reproducibility – was performed using deepTools (v3.5.3) through its bamPEFragmentSize, plotCoverage, and plotHeatmap modules[14].

##### *Multi-omic integration analyses*

EryP RNA-seq, pseudobulked scATAC-seq, and CUT&RUN were integrated at the gene level by calculating a gene score (scATAC-seq) or by taking the most prominent peak identified (CUT&RUN). $Log_2$FoldChange cutoffs of 0.25 were used to generate the venn diagrams in Figure 5I. Transcription factor enrichment was calculated using Fisher's exact test against reference hematopoietic transcription factor ChIP-seq obtained from the ENCODE Project or GEO.

#### Supplemental Figure Legends

**Supplemental Figure 1 HSPC scATAC-seq validation** (A) UMAP visualization of scATAC-seq cells from sorted *Stag2^WT^* and *Stag2^Δ^* HSPCs (n = 3 for both conditions) with all cell types annotated. (B) UMAP feature plots of marker genes of scATAC-seq cells filtered for the erythroid differentiation trajectory. (C) *Fli1* gene body accessibility over erythroid differentiation pseudotime.

**Supplemental Figure 2 Stag2 loss results in contraction of total MEPs and imbalances irreversible erythroid and megakaryocyte commitment** (A) Schematic diagram of bone marrow immunophenotyping for bipotent and unipotent erythroid and megakaryocytic progenitors. (B) Frequency of total MEPs (∑ PreMegE + MkP + EryP) from murine bone marrow after ACK lysis as determined by flow cytometry. (C) Schematic diagram of bone marrow immunophenotyping for irreversibly committed erythroid (top) and megakaryocyte cells (bottom). (D) Frequency of erythroid committed cells ($Ter119^{+}$; erythroblasts) from bone marrow of $Stag2^{WT}$ and $Stag2^{\Delta}$ mice after ACK lysis as determined by flow cytometry. (E) Representative flow cytometry plot identifying $Ter119^{+}$ cells stained with antibodies against CD11b, Gr-1, Kit, Ter119, and CD44. (F) Frequency of megakaryocytes ($Lineage^{-}Kit^{-}CD41^{+}$) from bone marrow after negative enrichment of lineage positive populations as determined by flow cytometry. (G) Representative flow cytometry plot identifying $CD41^{+}$ cells stained with antibodies against lineage markers (see supplemental table 1), Kit, CD41, CD55, CD105, and CD150. (H) Frequency of ProE from bone marrow after ACK lysis as determined by flow cytometry. Each point represents data from an individual mouse, bar graphs depict mean ± SD. * $p < 0.05$, ** $p < 0.01$, *** $p < 0.001$, **** $p < 0.0001$ for comparison of data from $Stag2^{WT}$ and $Stag2^{\Delta}$ assessed using unpaired two-tailed t test with Welch's correction ($\alpha = 0.05$) after using ROUT (Q=1%) method to remove outliers.

**Supplemental Figure 3 $Ter119^{+}$ scATAC-seq validation** A) UMAP visualization of scATAC-seq cells from sorted $Stag2^{WT}$ and $Stag2^{\Delta}$ $Ter119^{+}$ erythroblasts (n = 3 for both conditions) with standard morphologic nomenclature. (B) UMAP feature plots of marker genes of scATAC-seq cells. (C) Volcano plot representing statistical significance ($-\log_{10}FDR$) vs. magnitude of DNA accessibility changes ($\log_2 FoldChange$) from a comparison of pseudobulked $Stag2^{WT}$ and $Stag2^{\Delta}$ erythroblasts.

**Supplemental Figure 4 Mutual exclusivity of Gata1 cistrome intersections and integrated RNA/ATAC-seq plot** (A) Venn diagram depicting the number of upregulated and downregulated genes overlapping with Gata1 reference ChIP-seq as determined by CHEA3 in $Stag2^{WT}$ and $Stag2^{\Delta}$ EryPs. (B) Plot of magnitude of DNA accessibility changes vs. magnitude of RNA expression changes in $Stag2^{WT}$ and $Stag2^{\Delta}$ EryPs.

**Supplemental Figure 5 $Stag2^{\Delta}$ EryPs generate fewer cells** (A) Schematic diagram of liquid culture differentiation workflow. (B) Quantification of total number of cells present in each well after 48 hours of culture. (C) Representative brightfield microscopy of wells seeded with $Stag2^{WT}$ and $Stag2^{\Delta}$ EryPs after 48 hours. Images were obtained using a 10X/0.25 achromatic objective lens. The scale bar represents 50 μm. Each point represents data from an individual mouse, bar graphs depict mean ± SD. * $p < 0.05$, ** p

$< 0.01$, *** $p < 0.001$, **** $p < 0.0001$ for comparison of data from *Stag2*$^{WT}$ and *Stag2*$^{\Delta}$ assessed using unpaired two-tailed t test with Welch's correction ($\alpha = 0.05$) after using ROUT (Q=1%) method to remove outliers.

**Supplemental Figure 1**

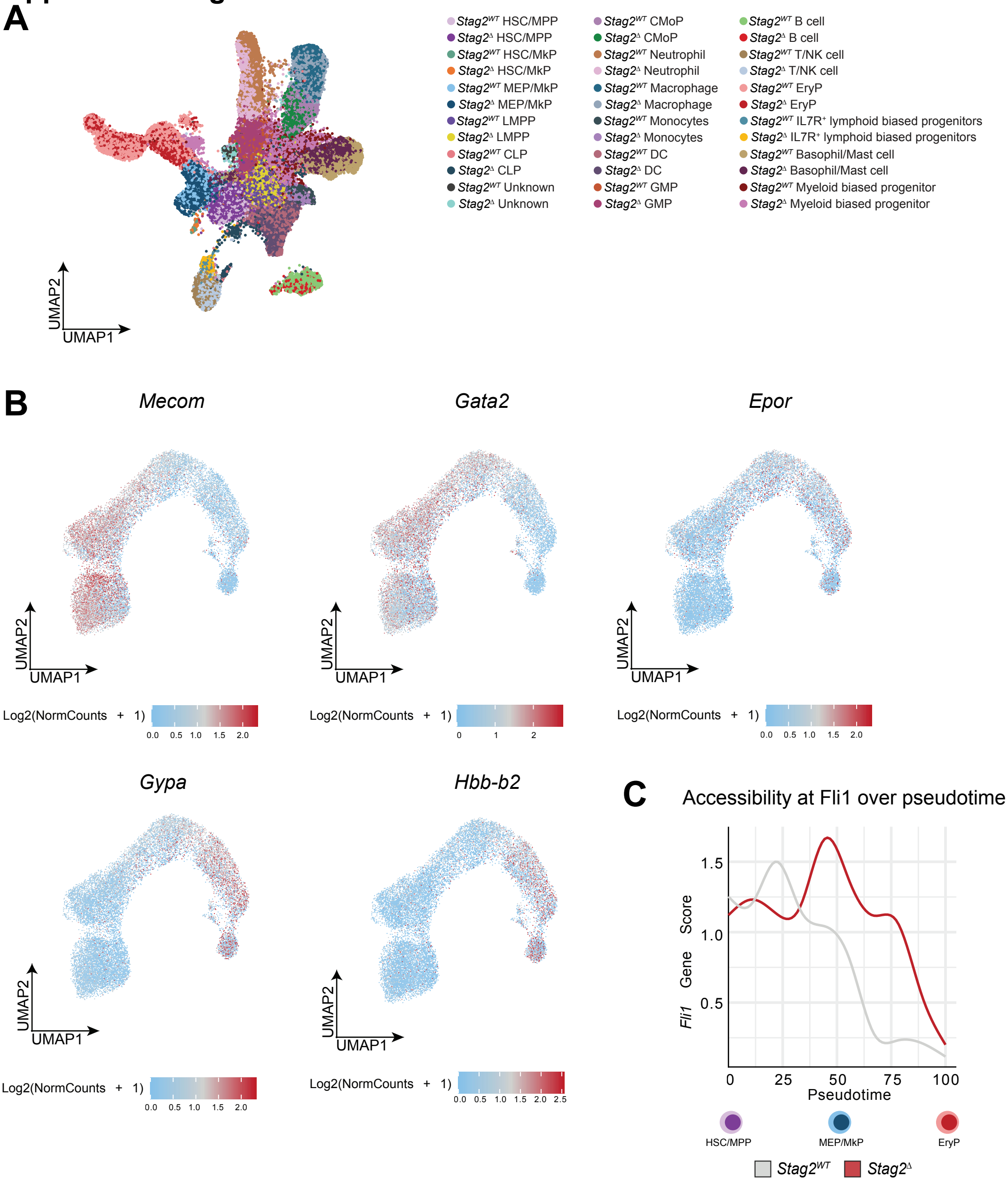

#### Supplemental Figure 2

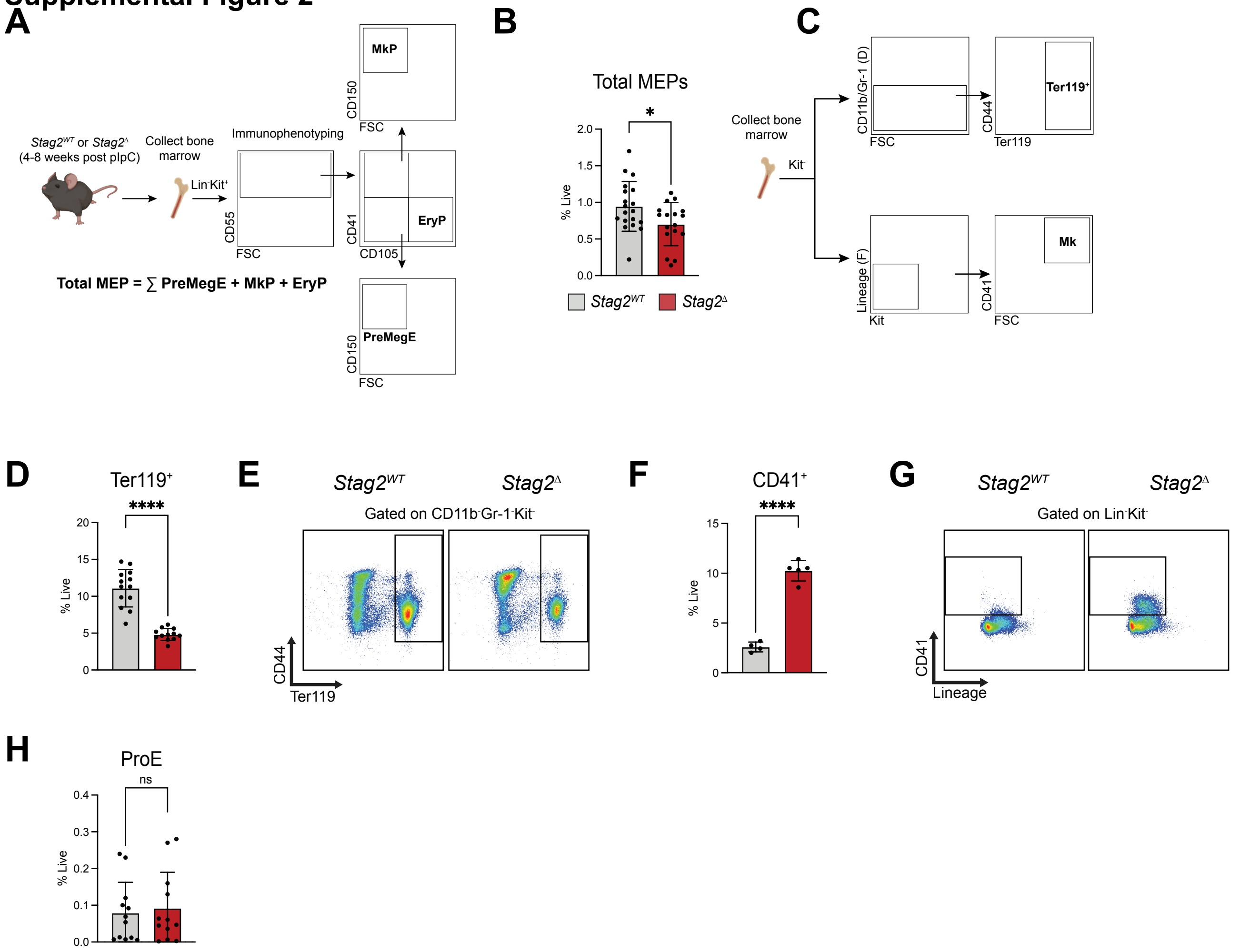

### Supplemental Figure 3

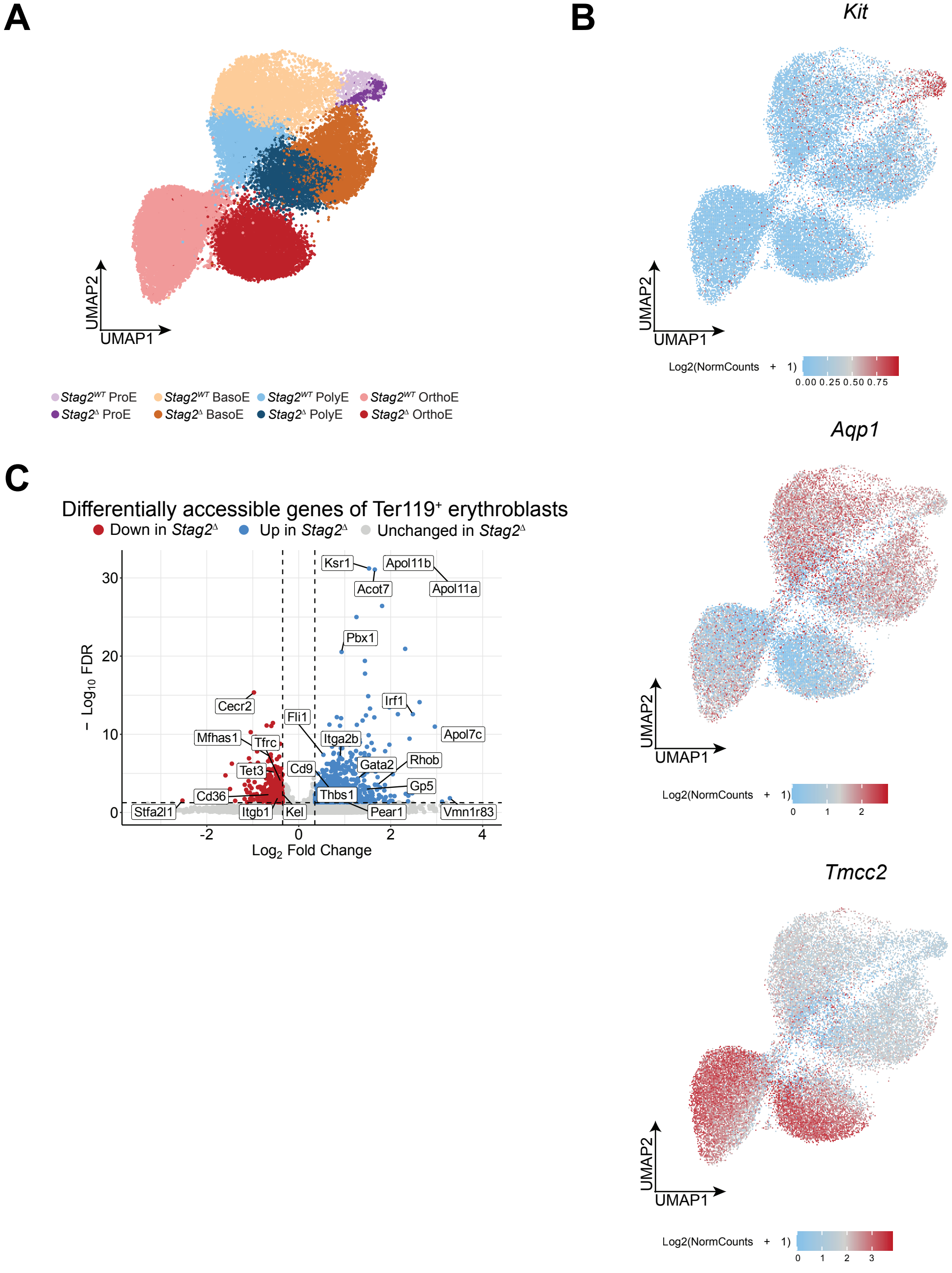

### Supplemental Figure 4

A

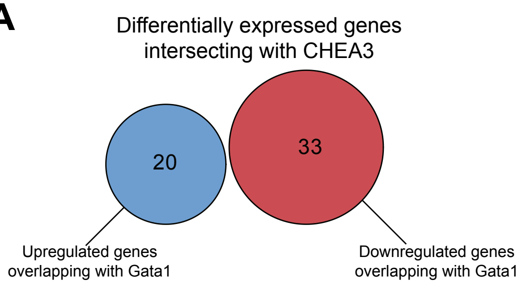


B

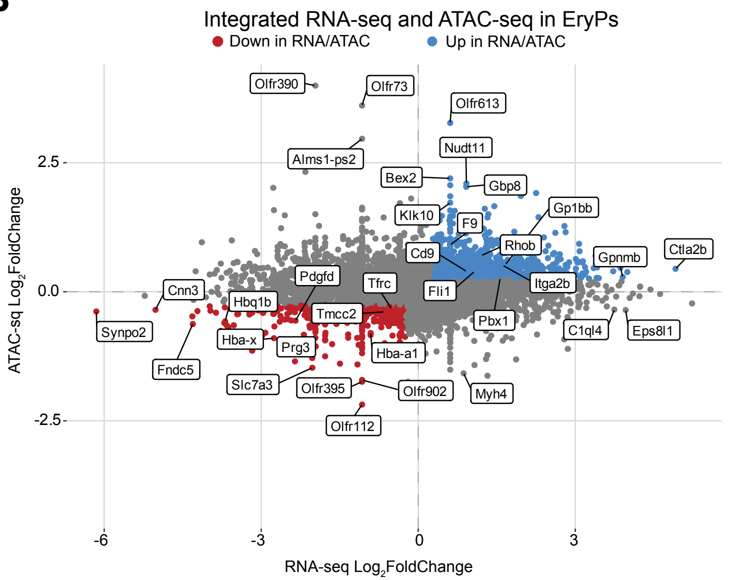

### Supplemental Figure 5

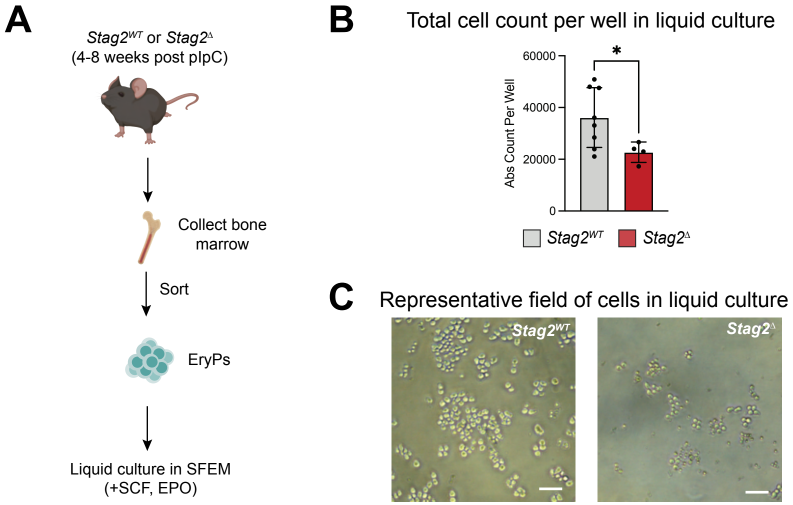
